# Role of zinc in growth, stress response and virulence gene expression of pathogenic Mucorale *Rhizopus arrhizus*

**DOI:** 10.1101/2025.10.13.681985

**Authors:** Rachna Singh, Anjna Kumari, Pavneet Kaur, Jasdeep Kaur

**Affiliations:** Department of Microbial Biotechnology, Panjab University, Chandigarh - 160014, India

**Keywords:** COVID-19, iron transport, mucormycosis, *Mucor*, mucoricin, *Rhizopus*, virulence, zinc

## Abstract

During COVID-19 pandemic, the cases of mucormycosis increased substantially, with rhino-orbito-cerebral form linked to uncontrolled diabetes being the predominant manifestation. Some clinical and epidemiological studies associated the usage of zinc supplements with the occurrence of COVID-19 associated mucormycosis, but experimental evidence remains limited. This study aimed to elucidate the impact of zinc enrichment on *Rhizopus arrhizus,* the predominant causative agent of mucormycosis. The effect of zinc supplementation (5 to 150 µM) on fungal growth, metabolic activity, antifungal susceptibility and biofilm formation, along with cell-wall (Congo red), oxidative (hydrogen peroxide) and osmotic (sodium chloride) stress was evaluated in RPMI-1640. Morphological changes were visualized by scanning electron microscopy. Gene expression was monitored by RNA sequencing. Zinc supplementation promoted *R. arrhizus* growth in a concentration-dependent manner, with the effect being particularly pronounced under less-favorable conditions, such as alkaline conditions resembling the nasal pH of diabetics. Exposure to zinc enhanced the metabolic activity, and partly alleviated the cell-wall and oxidative stress. Susceptibility to amphotericin B and posaconazole was unchanged. Zinc-supplemented cultures over-expressed multiple genes, notably those associated with respiratory electron transport chain, ergosterol biosynthesis, chitosan production, oxidative stress response, mucoricin, high-affinity iron permease, iron transport multi-copper oxidase and ferric-reductase-like protein. Zinc also augmented the growth/metabolism of several other *Mucorales* but not *Aspergillus fumigatus* and *Candida albicans*. The findings suggest that zinc supplementation supports mucoralean growth and enhances the expression of key virulence factors, implying that excessive intake of zinc supplements during COVID-19 pandemic likely contributed to the emergence of mucormycosis.

**IMPORTANCE:** Mucormycosis remains a highly devastating fungal infection that presented additional challenges during the COVID-19 pandemic. Several risk factors were implicated in the emergence of COVID-19 associated mucormycosis (CAM), including inappropriate corticosteroid usage and COVID-19 associated glycemic imbalance. The excessive intake of nutritional supplements, especially that of zinc, owing to self-medication and over-prescription, was also proposed as a plausible factor in the rise of CAM in many clinical/epidemiological studies. This study experimentally evaluated the potential impact of zinc enrichment on pathogenic *Mucorales.* The work demonstrates that zinc supplementation supports *Rhizopus arrhizus* growth, metabolism and stress response, and enhances the expression of key virulence factors, including mucoricin and iron transporters. Notably, the augmentation was specifically observed in pathogenic *Mucorales* but not *Aspergillus fumigatus* and *Candida albicans*. The work provides experimental evidence for the potential association of zinc over-availability with the occurrence of CAM, and strengthens evidence-based therapeutic practices in clinical settings.

## INTRODUCTION

The fungi belonging to the class Zygomycetes and the order *Mucorales* such as *Rhizopus, Rhizomucor, Lichtheimia*, *Mucor* and *Apophysomyces* are opportunistic pathogens that cause acute, angioinvasive infections associated with high morbidity and mortality (1–3). Mucormycosis is classified into distinct clinical entities, namely rhino-orbito-cerebral (ROC), pulmonary, cutaneous, gastrointestinal and disseminated types, with each being linked to specific risk factors (1–3). The disease is often community-acquired and transmitted through inhalation (ROC and pulmonary mucormycosis), ingestion (gastrointestinal mucormycosis) and occasionally via traumatic implantation (cutaneous mucormycosis) (1–3). The epidemiology of this disease varies substantially between the developed and the developing world. While ROC mucormycosis associated with uncontrolled diabetes is the predominant clinical manifestation in India, pulmonary mucormycosis associated with neutropenia and malignancies occurs more frequently in the developed countries (1–3). It is estimated that the prevalence of mucormycosis is nearly 70 times higher in India than the western world (1, 2). The problem was further aggravated by COVID-19, especially the second wave of this pandemic, with COVID-19 associated mucormycosis (CAM) being declared an epidemic in India (1).

Several risk factors were implicated in the emergence of CAM, including inappropriate corticosteroid usage, COVID-19 associated glycemic imbalance and immunosuppressive impact, microbiota dysbiosis, vascular endothelial injury, hypoxemia and dysregulated iron metabolism, along with increased expression of the mucoralean receptor GRP-78 in COVID-19 patients (1, 4–6). The excessive intake of nutritional supplements, especially that of zinc, owing to self-medication and over-prescription, was also proposed as a plausible factor in the rise of CAM (6–10). Zinc is an important micronutrient for growth and development of living beings, being crucial for gene expression, protein folding and intracellular signaling (11, 12). It is the second most abundant trace element in the human body, after iron, and its availability is tightly controlled by the mammalian host, as a part of nutritional immunity against microbial pathogens (11). Elevated levels of zinc can promote microbial growth (13–15), and some clinical and epidemiological studies reported an association of zinc intake with CAM (6–10). However, very few studies have experimentally determined the potential impact of zinc availability on mucoralean fungi (15–17). The present study therefore aimed to elucidate the effect of zinc enrichment on pathogenic *Mucorales,* with an emphasis on *Rhizopus arrhizus*, which is the predominant causative agent of mucormycosis worldwide.

## RESULTS

### Effect of zinc on *R. arrhizus* radial growth and metabolic activity

The effect on zinc on *R. arrhizus* growth was elucidated by radial growth assays in Roswell Park Memorial Institute medium (RPMI-1640) agar containing varying zinc concentrations (5 µM, 15 µM, 45 µM and 150 µM), and buffered to different pH (5.6, 7.0, 7.4 and 7.9). Addition of zinc to the culture medium impacted *R. arrhizus* growth in a pH and concentration-dependent manner (Table 1). At pH 5.6, *R. arrhizus* grew more profusely in the presence of zinc but the radial growth diameters in RPMI-1640 without or with zinc supplementation were similar (*P* > 0.05). As the pH of the culture medium increased to 7 and 7.4, the impact of zinc became more evident, and a significant, dose-dependent enhancement in fungal proliferation was noted with increasing zinc concentration (*P* < 0.05). *R. arrhizus* grew poorly at pH 7.9, and augmenting the culture medium with zinc substantially improved the fungal growth in a concentration-dependent manner (*P* < 0.05). Considering that pH 7.0 reasonably supported the fungal growth and depicted the dose-dependent alteration upon zinc supplementation, subsequent experiments were performed in RPMI-1640 at pH 7.0. Supplementation of the culture medium with zinc significantly increased the metabolic activity of *R. arrhizus*, indicated by a faster reduction of resazurin, a redox indicator dye that reflects the activity of electron transport chain (Figure 1; *P* < 0.05). The effect was evident even with 5 µM zinc (*P* < 0.05), and increasing the zinc level to 15 µM further hastened the dye reduction (*P* < 0.05). Whilst only 11% of the dye was reduced by *R. arrhizus* after 6 h of incubation in RPMI-1640 only, the percentage reduction increased to 51% upon addition of 5 µM zinc to the medium, and to nearly 74% with 15 µM zinc. No further enhancement was noted with 45 and 150 µM zinc, and the kinetics were nearly similar to those at 15 µM (Figure 1).

**Figure 1.**
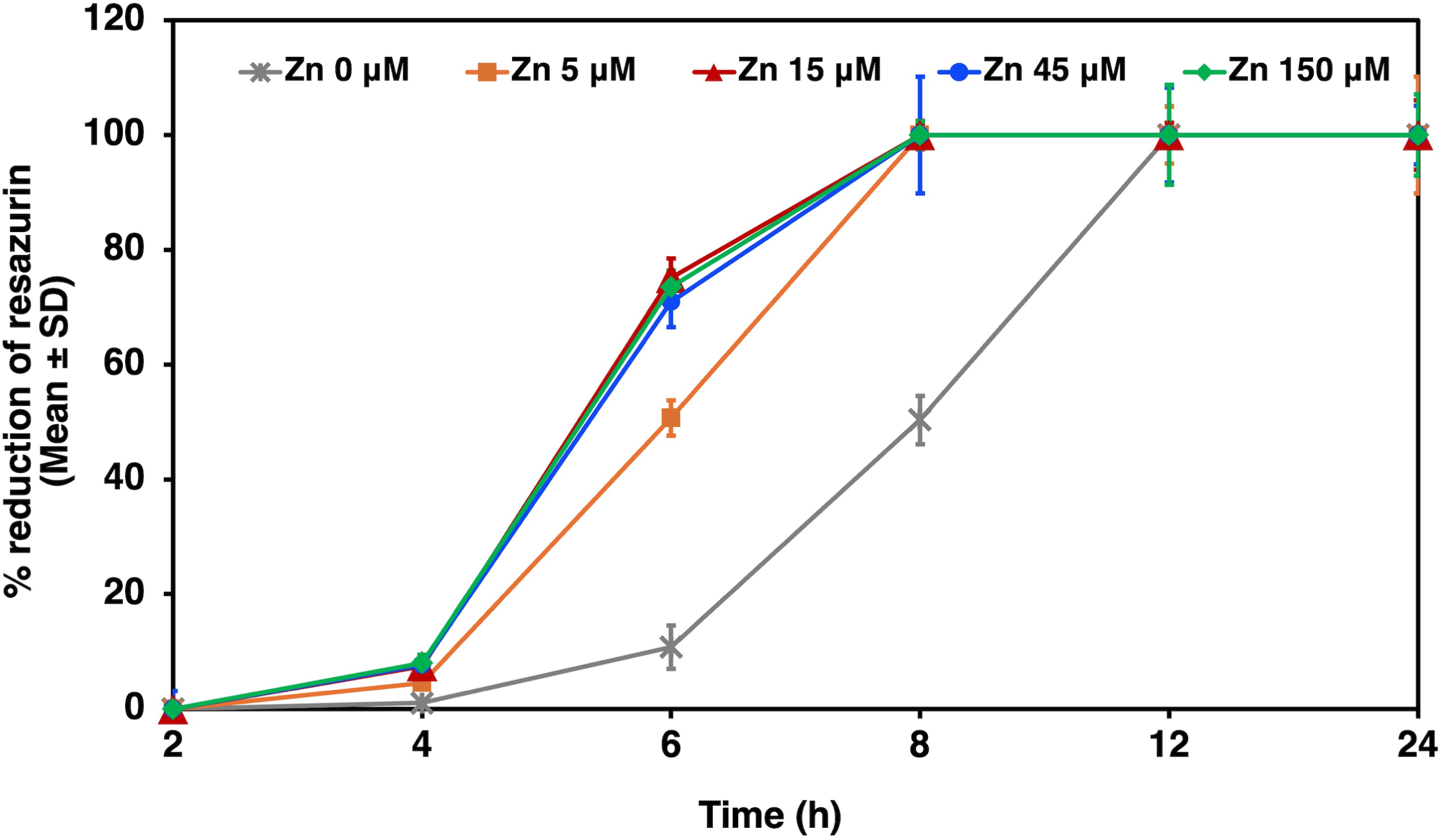
Metabolic activity of *Rhizopus arrhizus* NCCPF 710004 in RPMI-1640 (pH 7.0) supplemented with varying concentrations of zinc after incubation at 37°C for specified time intervals, as determined in terms of resazurin dye reduction (% reduction; Mean ± SD).

**Table 1.** Effect of varying concentrations of zinc on the radial growth of *Rhizopus arrhizus* NCCPF 710004 after incubation at 37°C in RPMI-1640 agar set to pH 5.6, 7.0, 7.4 and 7.9.

| | | Diameter of *R. arrhizus* radial growth (mm; Mean ± SD) | | | | |
| --- | --- | --- | --- | --- | --- | --- |
| pH | | Zn 0 μM | Zn 5 μM | Zn 15 μM | Zn 45 μM | Zn 150 μM |
| 5.6 | 24 h | 66.67 ± 0.47 | 70.5 ± 0.71 | 68.75 ± 0.35 | 70.75 ± 1.77 | 70 ± 0 |
| | 48 h | 85 ± 0<sup>#</sup> | 85 ± 0<sup>#</sup> | 85 ± 0<sup>#</sup> | 85 ± 0<sup>#</sup> | 85 ± 0<sup>#</sup> |
| 7.0 | 24 h | 21.5 ± 0.71 | 23.5 ± 0.71 | 27 ± 1.41 | 32 ± 1.41 | 43 ± 0.94 |
| | 48 h | 57 ± 0 | 60.5 ± 0.71 | 62.5 ± 0.71 | 76.5 ± 3.53 | 85 ± 0<sup>#</sup> |
| 7.4 | 24 h | 19.75 ± 1.06 | 20.5 ± 0.71 | 20.75 ± 0.35 | 23 ± 0.71 | 24.33 ± 0.47 |
| | 48 h | 48.75 ± 0.35 | 51.5 ± 0.71 | 52.25 ± 0.35 | 57.75 ± 0.35 | 59.5 ± 2.83 |
| 7.9 | 24 h | 8 ± 0 | 7.5 ± 0.71 | 7.75 ± 0.35 | 8.25 ± 0.35 | 10.5 ± 2.12 |
| | 48 h | 9.25 ± 0.35 | 9.5 ± 0.71 | 10 ± 0 | 10.5 ± 0 | 13.5 ± 1.4 |
| | 5 days | 14 ± 0.71 | 13.5 ± 0.71 | 18.75 ± 0.35 | 21.75 ± 2.47 | 29.67 ± 1.89 |

#, indicative of growth covering the entire petri plate

### Role of zinc in alleviating *R. arrhizus* stress

The impact of zinc enrichment in alleviating *R. arrhizus* stress was determined using Congo red, hydrogen peroxide and sodium chloride as cell-wall, oxidative and osmotic stressors respectively. The radial growth of *R. arrhizus* was markedly retarded by the cell-wall stressor Congo red (*P* < 0.05), and addition of zinc partially alleviated this effect in a dose-dependent manner (*P* < 0.05 at 48 h of growth; Figure 2). Zinc was also observed to partially ameliorate the growth-inhibitory impact of oxidative stress, particularly at 2 µM hydrogen peroxide (*P* < 0.05), as noted in terms of percentage spore germination (Table 2). At elevated levels of hydrogen peroxide (6 and 10 mM), the spore germination was fully inhibited regardless of zinc supplementation (*P* > 0.05; Table 2). Osmotic stress was not mitigated by this trace element, with the spore germination being considerably impeded upon exposure to sodium chloride (*P* < 0.05), independent of the presence or the level of the zinc in the growth medium (*P* > 0.05; Table 2).

**Figure 2.**
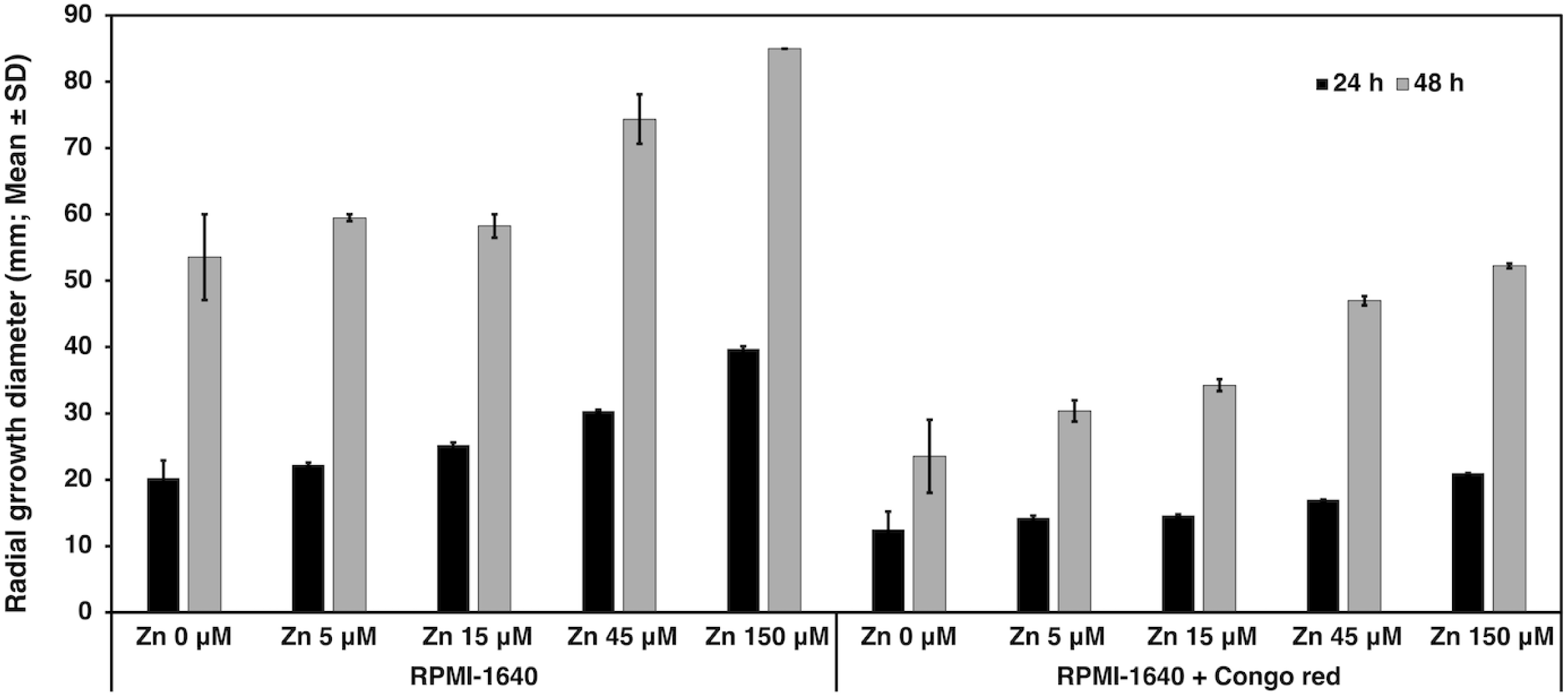
Radial growth diameters of *Rhizopus arrhizus* NCCPF 710004 (mm; Mean ± SD) upon exposure to the cell-wall stressor Congo red in RPMI-1640 agar (7.0) supplemented with varying concentrations of zinc after incubation at 37°C for 24 h and 48 h.

**Table 2.** Spore germination of *Rhizopus arrhizus* NCCPF 710004 upon exposure to oxidative stressor (hydrogen peroxide; H_2_O_2_) and osmotic stressor (sodium chloride; NaCl) after incubation in RPMI-1640 (pH 7.0) supplemented with varying concentrations of zinc at 37°C for 6 h.

| *R. arrhizus* spore germination (%; Mean ± SD) |  |  |  |  |  |
| --- | --- | --- | --- | --- | --- |
|  | Zn 0 µM | Zn 5 µM | Zn 15 µM | Zn 45 µM | Zn 150 µM |
| **RPMI-1640** | 77.02 ± 4.80 | 71.23 ± 2.57 | 76.06 ± 7.57 | 76.06 ± 6.23 | 74.20 ± 6.19 |
| + 2 µM H<sub>2</sub>O<sub>2</sub> | 46.86 ± 1.15 | 59.23 ± 2.11 | 62.36 ± 0.94 | 62.75 ± 3.61 | 64.71 ± 3.70 |
| + 6 mM H<sub>2</sub>O<sub>2</sub> | 0 | 0 | 0 | 0 | 0 |
| + 10 mM H<sub>2</sub>O<sub>2</sub> | 0 | 0 | 0 | 0 | 0 |
| + 0.145 M NaCl | 47.24 ± 3.34 | 49.21 ± 1.38 | 50.62 ± 0.26 | 48.47 ± 3.02 | 54.74 ± 4.54 |
| + 0.435 M NaCl | 0 | 0 | 0 | 0 | 0 |
| + 1.45 M NaCl | 0 | 0 | 0 | 0 | 0 |

### Antifungal drug susceptibility

The susceptibility of *R. arrhizus* to amphotericin B and posaconazole, with and without zinc supplementation, was determined by broth microdilution method. The minimum inhibitory concentrations of amphotericin B (0.125 µg/ml) and posaconazole (1 µg/ml) against *R. arrhizus* were unchanged by zinc levels in the culture medium.

### Effect of zinc on biofilm-forming capacity

*R. arrhizus* biofilm-forming capacity was found to decrease significantly (*P* < 0.05) upon supplementation of the growth medium with zinc, as noted microscopically and quantified in terms of A_490_ of the bound safranin (Figure 3). Furthermore, the mycelial aggregation exhibited an inverse relation with zinc levels, with densely-aggregated hyphae being noted at 0 and 5 µM zinc, and loose aggregates observed at 5 to 150 µM zinc in a dose-dependent fashion (Figure 3).

**Figure 3.**
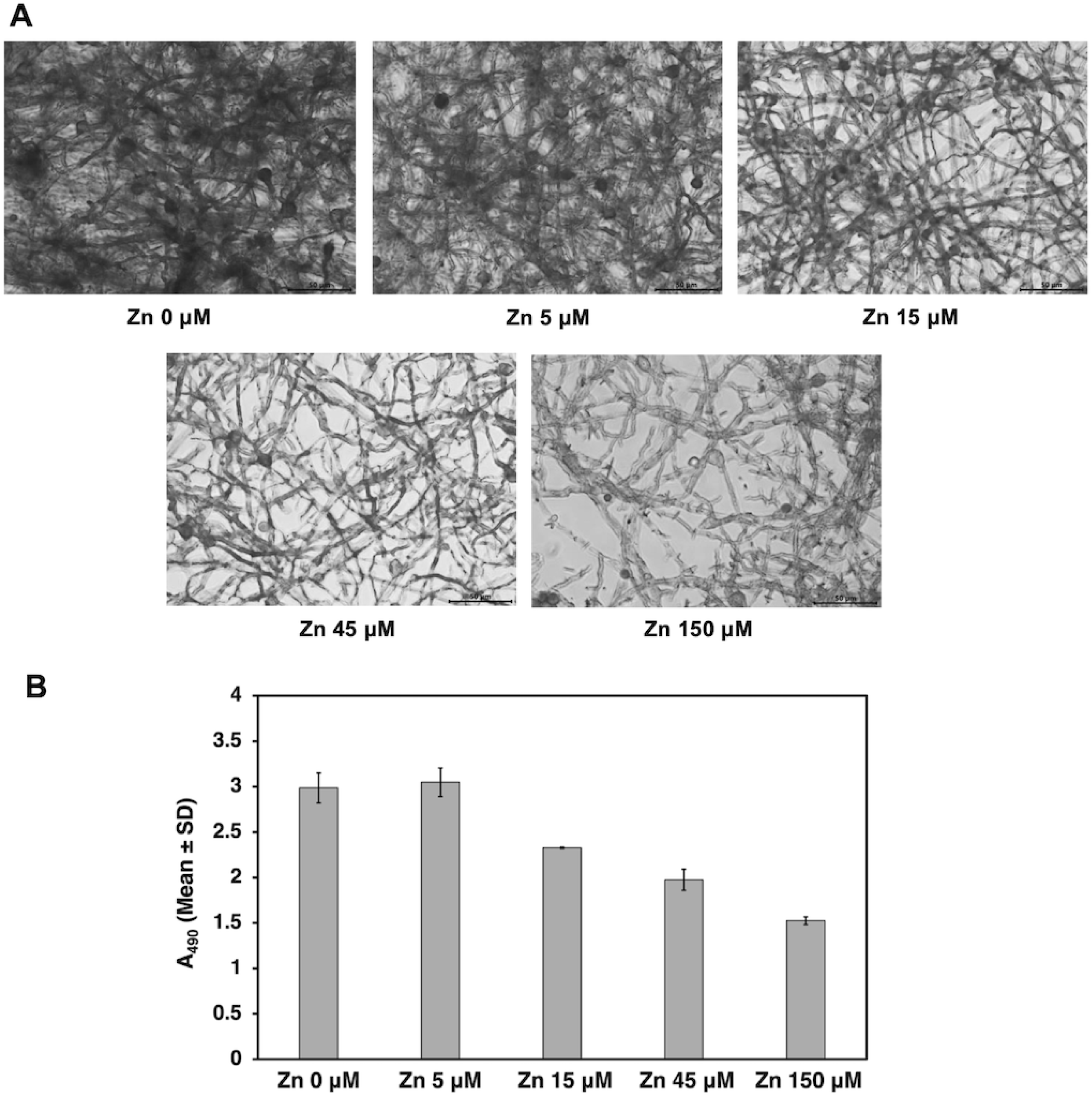
Biofilm-forming capacity of *Rhizopus arrhizus* NCCPF 710004 in RPMI-1640 (pH 7.0) supplemented with varying concentrations of zinc after incubation at 37°C for 24 h, as noted by inverted microscopy (A) and quantified by measuring the absorbance of bound safranin (B; A_490_; Mean ± SD). Magnification, 400x; Scale bar 50 µm.

### Liquid culture phenotype with zinc supplementation

The macro-morphology of *R. arrhizus* liquid cultures changed upon zinc enrichment, with cultures grown in RPMI-1640 only forming distinct pellets compared to loose aggregates and dispersed mycelia seen in zinc-supplemented media (Figure 4A). Field-emission scanning electron microscopy (FE-SEM) corroborated these findings and further demonstrated that the hyphae cultured in zinc-supplemented media were comparatively thinner and had more pronounced surface undulations than those in RPMI-1640 only (Figure 4B). Zinc also prolonged the growth of *R. arrhizus* in broth cultures, delaying the onset of decline phase, as noted in mycelial dry weight experiments (Figure 4C). *R. arrhizus* grown in RPMI-1640 without zinc supplementation reached the peak mycelial dry weight within 24 h of incubation, followed by a decline in mycelial biomass day 2 onwards. In contrast, when cultured in the presence of 15 µM and 45 µM zinc, the peak mycelial dry weights were achieved at 2 and 4 days of incubation respectively, with the decline phase beginning at 4 and 6 days, respectively (Figure 4C).

**Figure 4.**
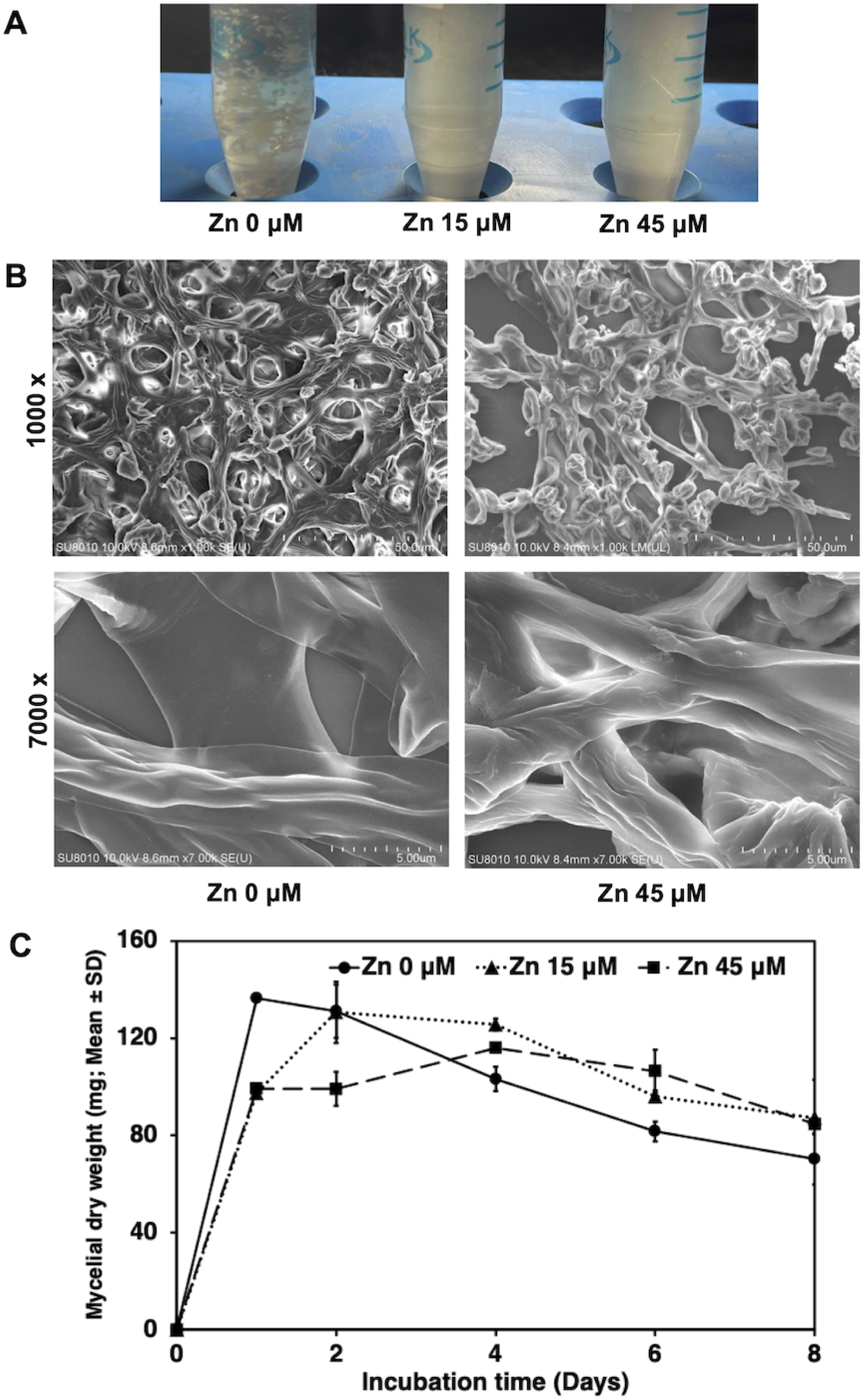
Liquid culture phenotype of *Rhizopus arrhizus* NCCPF 710004 in RPMI-1640 (pH 7.0) supplemented with zinc. **A,** Representative images demonstrating the macro-morphology of *R. arrhizus* cultures without and with zinc supplementation after incubation at 37°C for 6 h. **B,** Field-emission scanning electron micrographs of *R. arrhizus* in culture medium without and with zinc supplementation. The images were taken at a magnification of 1000× and 7,000×. Scale bar, 50 μm and 5 μm for images at 1000× and 7,000× respectively. **C,** Mycelial dry weight of *R. arrhizus* (mg; Mean ± SD) achieved without and with zinc supplementation after incubation at 37°C for specified time intervals.

### RNA sequencing of zinc-supplemented *R. arrhizus* cultures

Enrichment of the culture medium with zinc led to an up-regulation of 273 genes and down-regulation of 265 genes in *R. arrhizus* (Table 3, Figure 5 and supplementary data S1 and S2). A variety of genes involved in amino-acid metabolism (in terms of functional enrichment and biological processes); ribosome biogenesis, translation and protein folding (in terms of cellular component); along with nucleic-acid binding and translation-regulator activity (in terms of molecular function) were over-expressed (Figure 5). Notably, zinc-supplemented cultures exhibited an increased expression of genes involved in ergosterol biosynthesis (Table 3). The expression of cell-wall enzymes was modulated, with chitinase being down-regulated, and chitin deacetylase being up-regulated (Table 3). The oxidative stress response genes, including superoxide dismutase (*sod*1) and glutathione S-transferase were also over-expressed upon zinc addition (Table 3). Additionally, many of the genes involved in respiratory electron transport chain were up-regulated (Table 3). Amongst the virulence factors, the genes involved in iron transport, such as the high-affinity iron permease (*ftr*1p), the iron transport multi-copper oxidase (*fet*) and ferric-reductase-like protein (*fre*), as well as the mucoralean toxin mucoricin were up-regulated with zinc enrichment (Table 3). The expression of mucoralean invasin *cot*H genes, siderophore rhizoferrin, ADP-ribosylation factor, ADP-ribosylation factor-like protein 1, rhizopuspepsin protease, RNA silencing genes (RNA dependent RNA polymerase and Dicer-like protein 1) and deferoxamine uptake proteins (Fob1 and Fob2), and regulatory pathways including calcineurin regulatory subunit B, calcium-calmodulin dependent protein kinases, cAMP-dependent protein kinase A, mitogen activated protein kinases and Ras-related GTPases (*rho*, *rac* and *cdc*42) was unaffected (Table 3). However, the expression of *rab*6A, and many of the regulators of Ras-related GTPases (*rho*, *rac* and *cdc*42) (Table 3) was altered with zinc exposure, suggesting a potential modulation of actin cytoskeleton and/or cell morphogenesis (Table 3). Zinc transporters (ZIP), genes involved in sulfur metabolism (*cys*H), translation elongation factor (EF3 and TEF3), along with aspartic-type (SAP) and serine-type proteases (subtilase) were amongst the key genes down-regulated by zinc exposure (Table 3).

**Figure 5.**
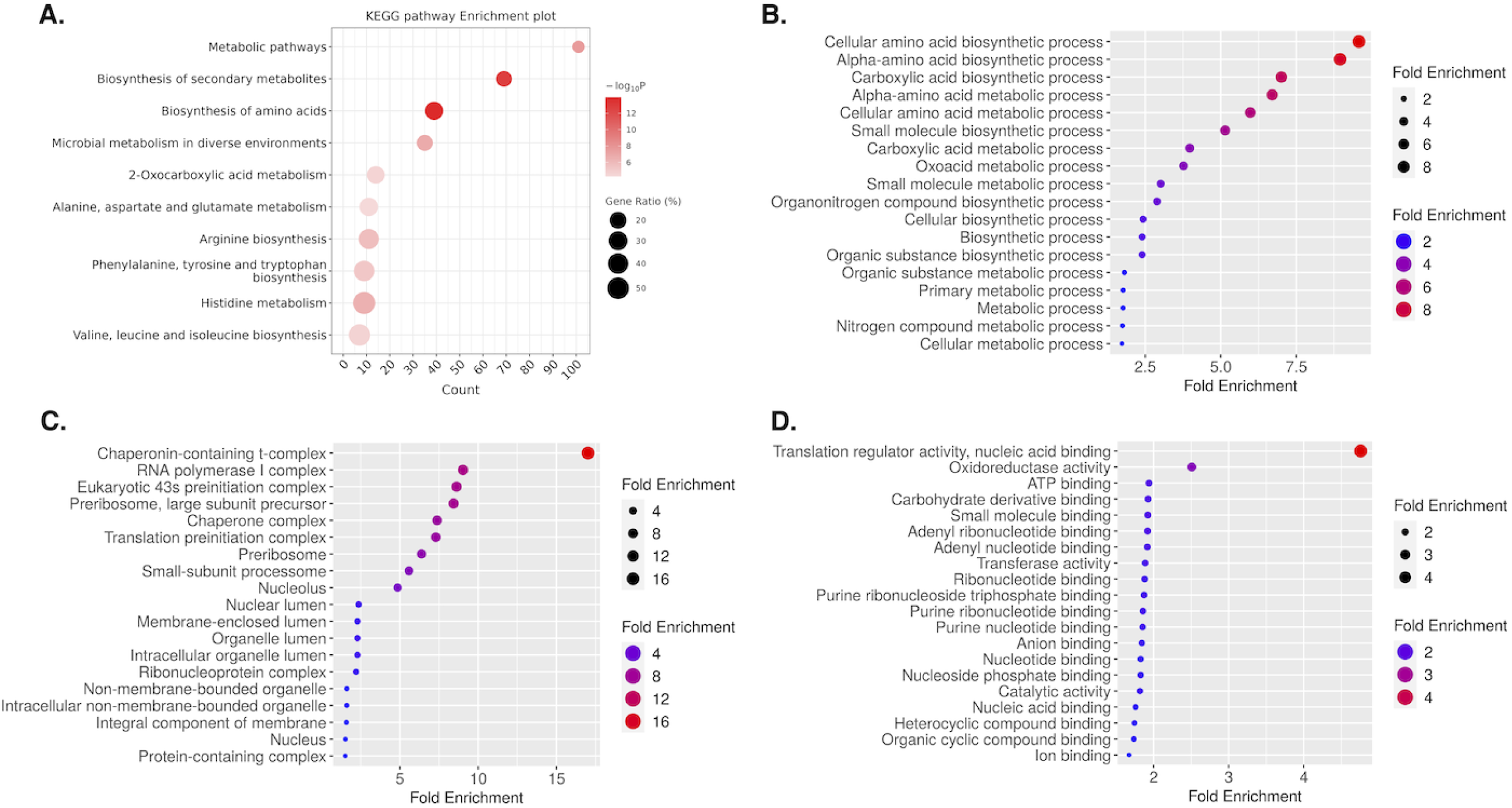
Gene expression of *Rhizopus arrhizus* NCCPF 710004 after culturing in RPMI-1640 (pH 7.0) supplemented with 45 µM zinc compared with the RPMI-1640 only control, as determined by RNA transcriptome sequencing. **A-D,** Functional enrichment showing the top-most significant (FDR<0.05) categories of KEGG terms (A), biological processes (B), cellular component (C) and molecular function (D) GO terms based on up and down regulated genes of the test setup (with Zinc 45 µM) vs. the control. The top GO terms are plotted in terms of fold change. The size of the dots represents the number of genes in the significant differential expression gene list associated with the GO term and the color of the dots represent the fold enrichment.

**Table 3.**
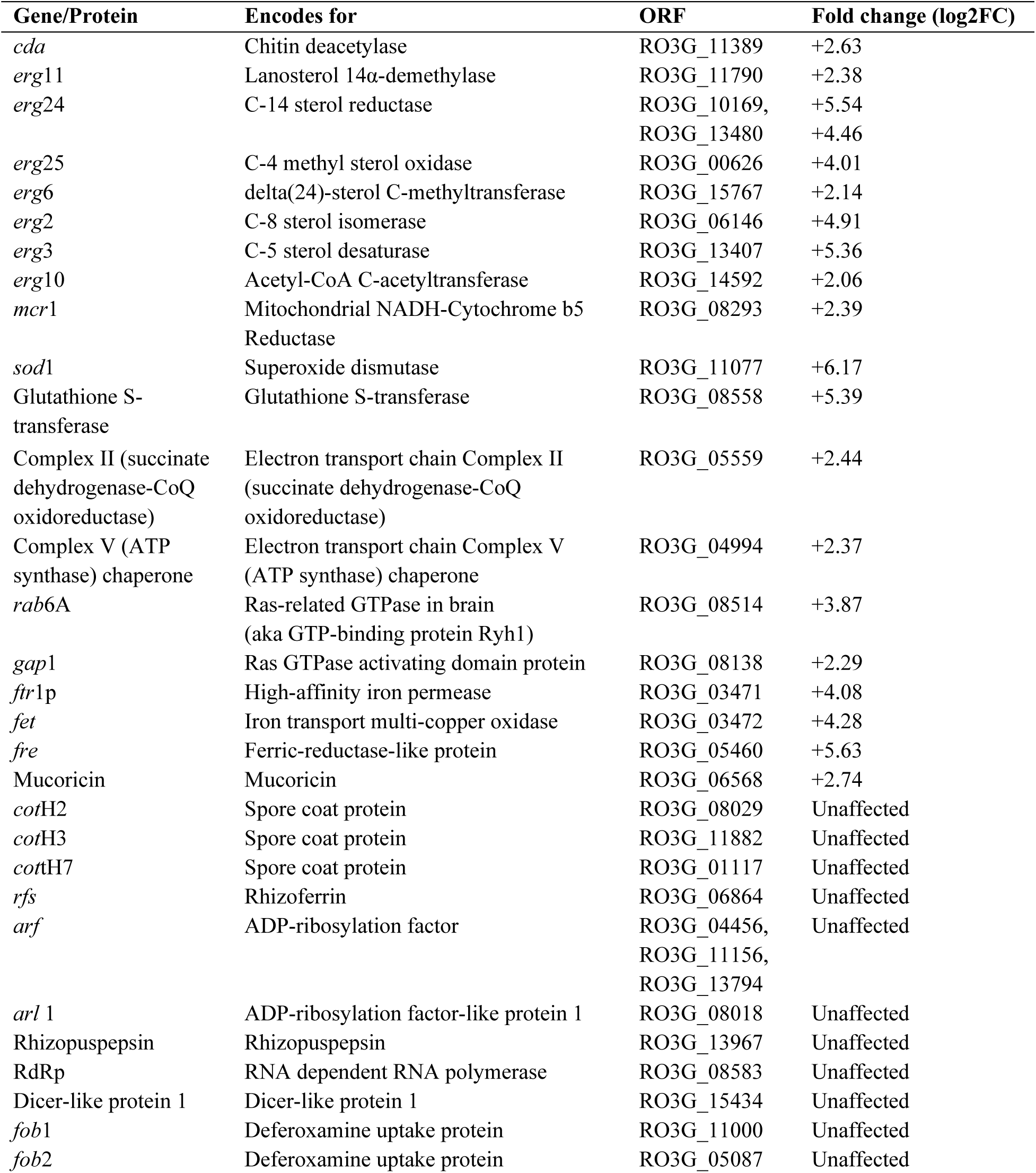
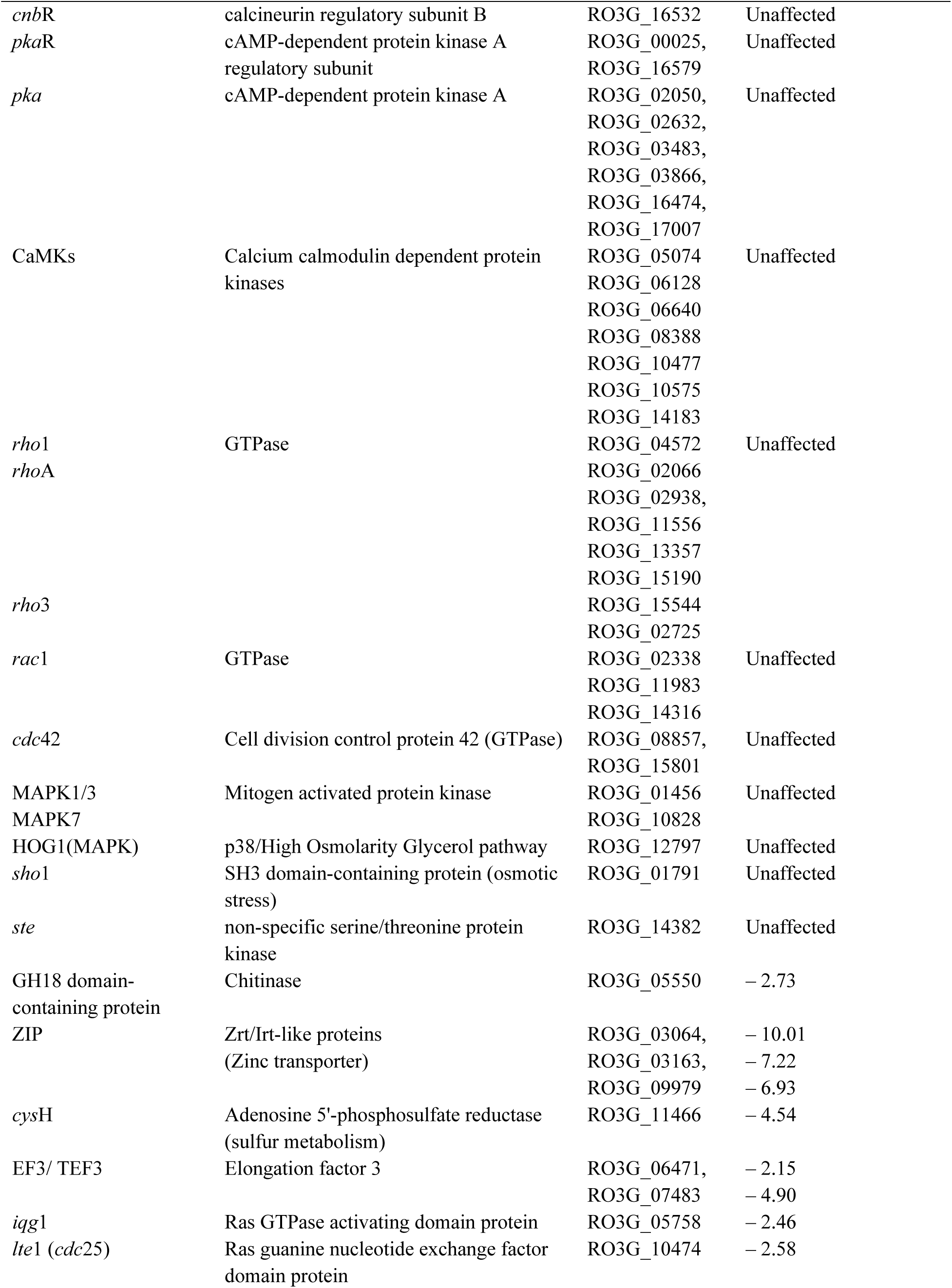
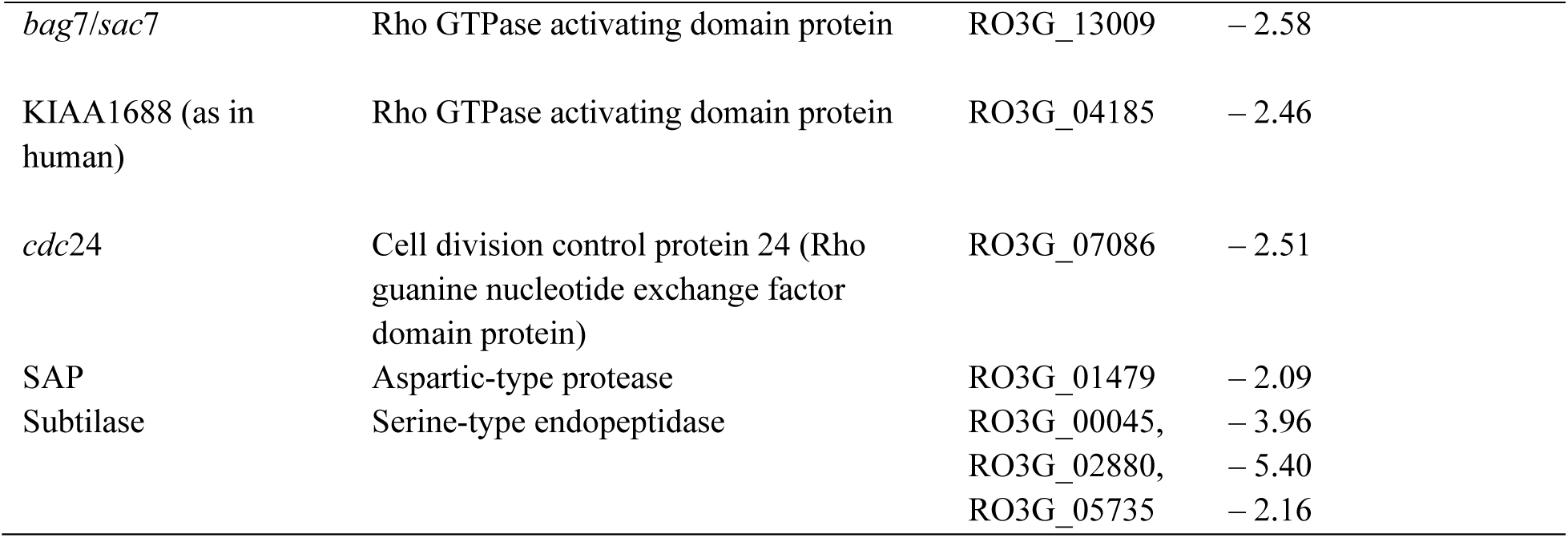
Impact of zinc enrichment on the gene expression of *Rhizopus arrhizus* NCCPF 710004 as determined by RNA transcriptome sequencing. The table depicts the expression of key *R. arrhizus* genes/proteins that were majorly influenced by zinc exposure, and those that were unaffected, with an emphasis cell growth and virulence-related genes.

### Effect of zinc on other fungal strains

The effect of zinc was then evaluated on growth and metabolic activity of additional mucoralean species/strains (*R. arrhizus* NCCPF 710003*, Rhizomucor pusillus* NCCPF 720004, *Lichtheimia corymbifera* NCCPF 700002*, Cunninghamella bertholletiae* MTCC 8869, *Mucor indicus* MTCC 3318 and *Mucor irregularis* MTCC 10708), along with two other common invasive fungi *Aspergillus fumigatus* NCCPF 770372 and *Candida albicans* ATCC 90028. Radial growth based assays were employed for assessing the growth of filamentous fungal strains, and liquid culture growth curves were used for *C. albicans*. The growth of *A. fumigatus* and *C. albicans* was unaffected by addition of zinc to the culture medium (Figure 6A and 7A-D). Their metabolic activity also remained unchanged, with a decline noted at high zinc concentrations likely due to the toxicity associated with this heavy metal (Figure 6B and 7E). On the contrary, all the mucoralean strains tested showed significantly increased radial growth diameters in zinc-supplemented media, especially at pH 7.9 (*P* < 0.05; Figure 8A), except *L. corymbifera*. A qualitative enhanced (denser) radial growth was also observed for *L. corymbifera* although the growth diameters were unchanged (without zinc, 68 ± 1.14; with 150 µM zinc, 69 ± 0.82 mm). Similarly, the metabolic activity of majority of the *Mucorales* was substantially fastened upon zinc supplementation (*P* < 0.05) (Figure 8B). For *R. pusillus*, rezasurin was fully reduced even in the control set ups after 24 h of incubation, while shorter incubation periods resulted in negligible reduction.

**Figure 6.**
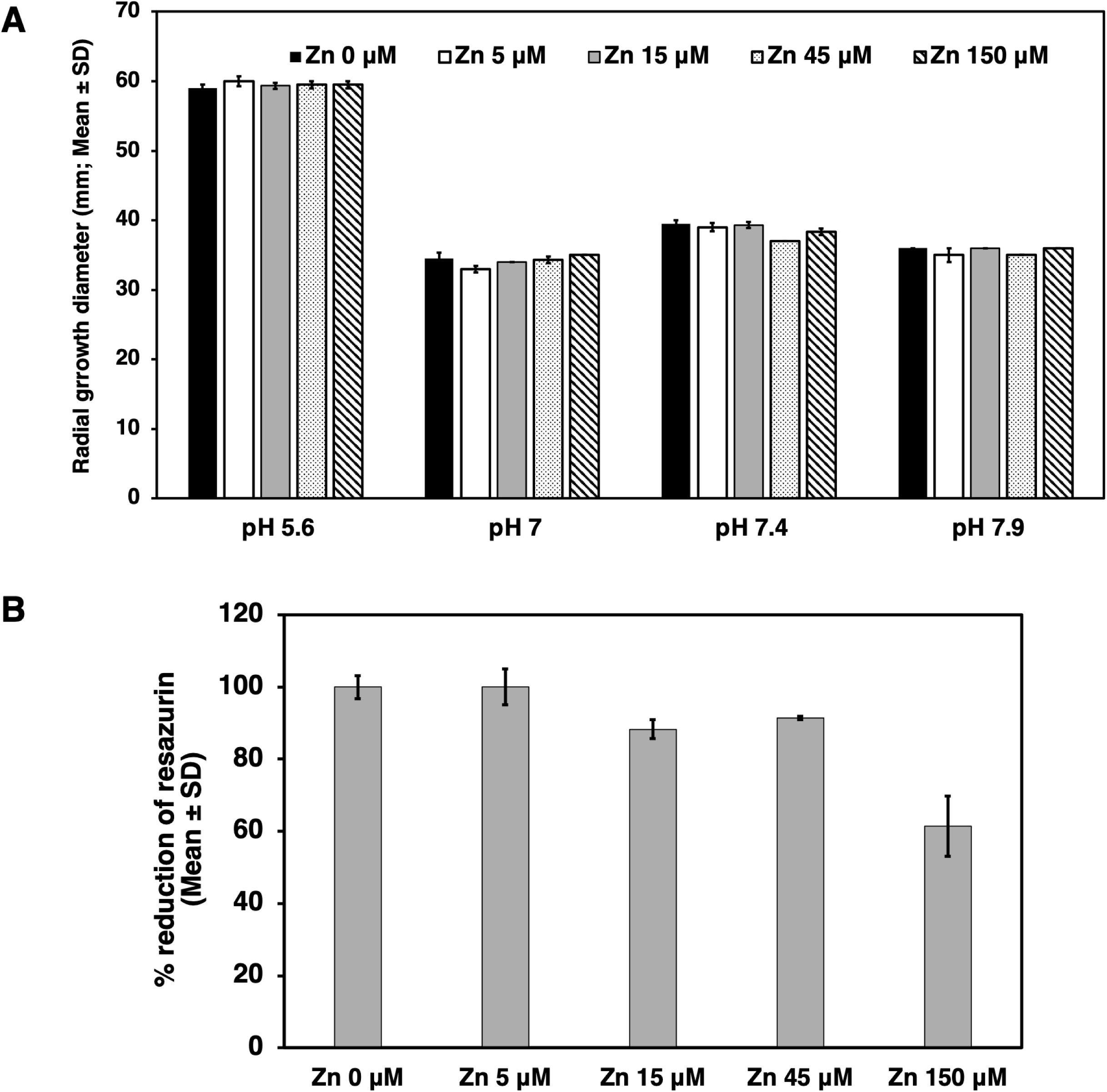
**A**, Radial growth of *Aspergillus fumigatus* NCCPF 770372 in RPMI-1640 agar at pH 5.6, 7.0, 7.4 and 7.9 after incubation at 37°C for 48 h, without and with zinc supplementation. **B**, Metabolic activity in RPMI-1640 (pH 7.0) supplemented with varying levels of zinc after incubation at 37°C, as determined in terms of resazurin dye reduction (% reduction; Mean ± SD).

**Figure 7.**
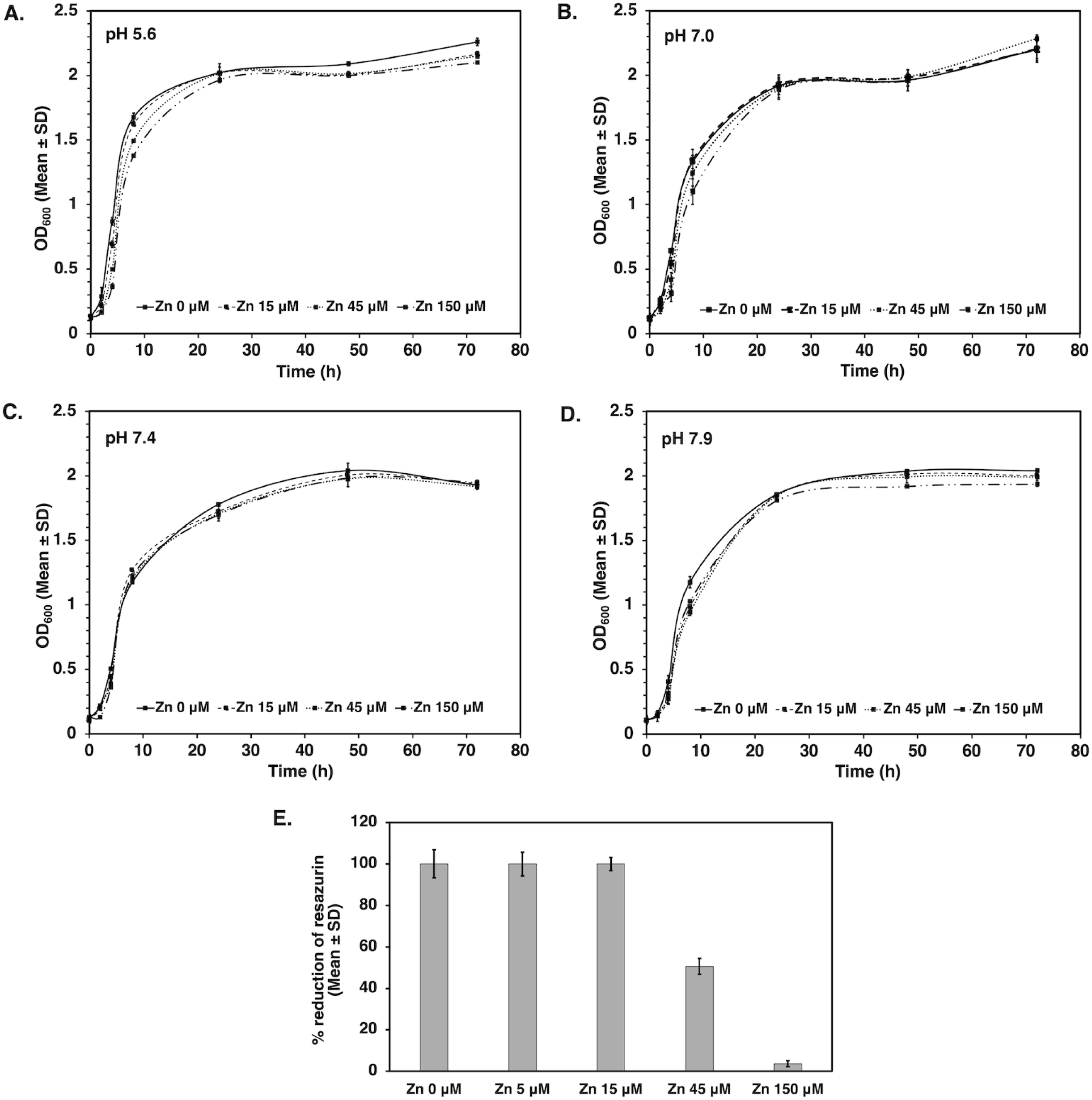
**A-D,** Growth curves of *Candida albicans* ATCC 90028 in RPMI-1640 at pH 5.6, 7.0, 7.4 and 7.9 after incubation at 37°C for specified time intervals, without and with zinc supplementation. **E,** Metabolic activity in RPMI-1640 (pH 7.0) supplemented with varying levels of zinc after incubation at 37°C, as determined in terms of resazurin dye reduction (% reduction; Mean ± SD).

**Figure 8.**
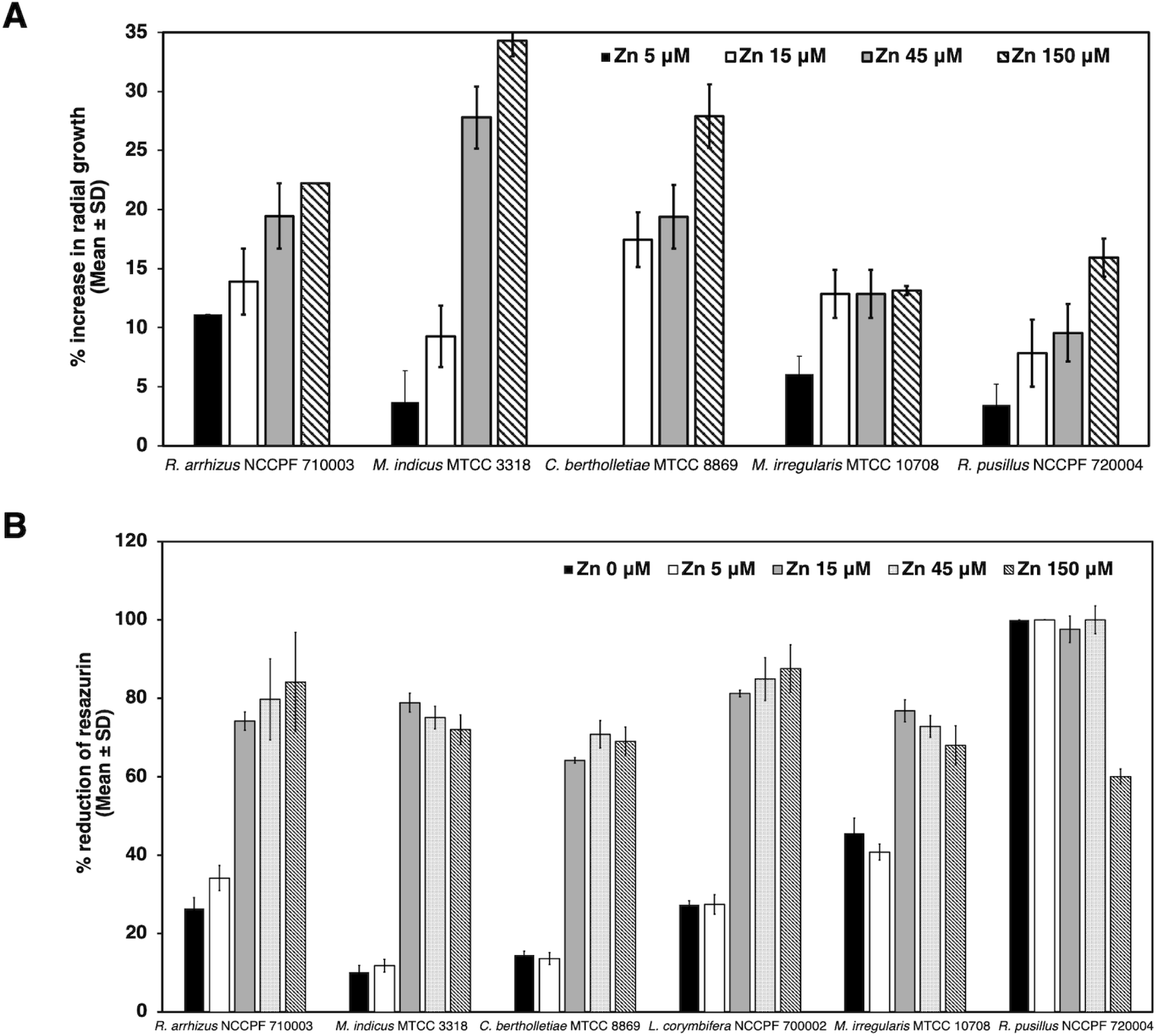
**A,** Effect of varying concentrations of zinc on the radial growth of different mucoralean species (mm; Mean ± SD) after incubation at 37°C (30 ℃ for *Mucor* spp.) in RPMI-1640 agar for 48 h. **B,** Metabolic activity of various mucoralean species in RPMI-1640 supplemented with varying levels of zinc after incubation at 37°C (30 ℃ for *Mucor* spp.), as determined in terms of resazurin dye reduction (% reduction; Mean ± SD).

## DISCUSSION

The devastating consequences of COVID-19 pandemic were worsened by the rising burden of secondary infections, including invasive mold infections caused by *Aspergillus* spp., *Candida* spp., and *Mucorales* (1). An unprecedented surge in mucormycosis cases was particularly astonishing. Several reasons were proposed to explain the occurrence of CAM, with COVID-19 associated glycemic imbalance and inappropriate glucocorticoid usage being the major ones (1). In addition, the increased prophylactic and therapeutic usage of zinc supplements as a measure against COVID-19 was noted as a risk factor for CAM in some case-control studies (6–10). Zinc was prescribed for management of COVID-19 due to its antioxidant, anti-inflammatory, immune-supporting, and potential antiviral effects (12, 18). Self-medication with zinc supplements was also common during that period. However, being an essential trace element for all life-forms, zinc overload may increase susceptibility to microbial infections (19). It is known to impact microbial growth (13–15) and zinc chelators exhibit antimicrobial activity (17). Muthu *et al.* have documented that *R. arrhizus* grows more profusely in zinc-enriched media, with some isolates showing higher viable counts (16). However, a detailed mechanistic investigation into the influence of zinc augmentation on mucoralean fungi is largely lacking. This study assessed the impact of zinc on growth, stress response and virulence gene expression of the pathogenic mucorale *R. arrhizus*.

The effect of zinc on *R. arrhizus* was tested by supplementing RPMI-1640 with a range of zinc concentrations (5 to 150 µM) to exemplify zinc deficit and overload, and zinc levels across different body sites, with 15 µM depicting the normal serum level and 150 µM being the total concentration in brain (20). The culture medium RPMI-1640 lacks zinc, and the experiments were conducted in type I water; however, no specific attempt was made to eliminate any traces of zinc sourced in the experimental setups through glassware and reagents. The growth of *R. arrhizus* was evaluated in culture media set to pH 5.6, 7, 7.4 and 7.9, representing the physiological pH of healthy and diabetic individuals in the nasal cavity (6.0 and 7.9 respectively) and the serum (7.3-7.4 and 6.6-7.3 respectively) (21, 22). Radial growth measurement assays revealed that zinc supplementation supported the growth of *R. arrhizus per se,* and particularly under less favorable pH conditions (neutral and alkaline vs. acidic), with higher levels of zinc (45 and 150 µM) leading to a more pronounced growth enhancement. Considering that the nasal pH in diabetic patients is alkaline (21) the findings suggest that elevated zinc levels are conducive for *R. arrhizus* in the context of ROC mucormycosis, which is associated with uncontrolled diabetes and acquired by nasal inhalation (1, 2). In healthy individuals, the fungal growth and invasion would be controlled by the fully functional immune system and the low-level expression of the mucoralean host receptor GRP-78 (1, 2). The elevated fungal proliferation associated with zinc enrichment may, however, overwhelm the capacity of phagocytic cells to respond effectively in healthy individuals as well (16, 23).

Enriching the culture medium with zinc was found to substantially fasten *R. arrhizus* metabolism. An earlier study also reported that zinc increases the efficiency of energy utilization, leading to rapid and abundant growth, and a more complete destruction of glucose (15). Furthermore, the metal prolonged *R. arrhizus* growth in liquid cultures and delayed the onset of decline phase, plausibly due to the growth morphotype observed with zinc addition. In the presence of zinc, *R. arrhizus* cultures formed loose aggregates and/or dispersed hyphae, which likely supported enhanced mass transfer and nutrient availability (24), unlike the compact pellets observed without zinc. Consistent with these findings, the biofilm-forming ability was observed to be inversely related to zinc levels, as the zinc-induced spatial dispersion would inhibit the formation of compact aggregates that are necessary for biofilm development.

Zinc constitutes the catalytic and structural site of several enzymes, and a huge number of transcription factors contain zinc binding motifs (11). Given the crucial role of this heavy metal in the synthesis, stabilization and regulation of proteins and nucleic acids (14, 25), zinc-enriched *R. arrhizus* cultures were found to over-express a wide range of genes. In line with the impact of zinc on growth and metabolism of *R. arrhizus*, the expression of many of the genes involved in cellular respiration increased with zinc exposure. The biosynthetic pathway for ergosterol, the fungal cell membrane sterol, was also found to be up-regulated following zinc supplementation, as also reported earlier for *C. albicans* (25). Additionally, zinc enrichment led to the down-regulation of chitinase and up-regulation of chitin deacetylase, suggesting an increase in the levels of chitosan, which is a distinct component of the mucoralean cell wall (26). This corroborated with the findings of cell-wall stress assay, wherein zinc partially alleviated *R. arrhizus* growth during Congo red induced stress. FE-SEM demonstrated that the hyphal surface of zinc-supplemented cultures exhibited increased surface undulations. This is likely attributed to the observed over-production of cell-wall (chitosan) and cell-membrane (ergosterol) constituents, and plausibly the cytoskeletal rearrangements, as the expression of many of the regulators of Ras-related GTPases was altered with zinc exposure. A previous study has reported that zinc alters the cell morphology, increases chitin deposition and thickens the cell wall of an ericoid ascomycete fungus (27). Such structural alterations are known to influence the fungal pathogenicity (28). For instance, chitosan levels have been directly linked to germling adherence, fungal virulence and enhanced protection from host immune response in pathogenic fungi (29, 30). Likewise, ergosterol biosynthetic pathway modulates the growth, membrane stability, sporulation, virulence and antifungal susceptibility in *Mucor* sp. (31).

Zinc partly mitigated the inhibitory impact of 2 µM hydrogen peroxide, a concentration corresponding to the physiological levels seen in phagocytic cells (32). The higher concentration of hydrogen peroxide (6 mM) is known to induce apoptosis in *R. arrhizus* (33), and was found to be growth inhibitory in the present study as well, independent of zinc supplementation. Additionally, the expression of many genes involved in oxidative stress, including *sod*1 and glutathione S-transferase was enhanced by zinc enrichment. These data correlate with the pivotal role of zinc in eukaryotic antioxidant defense system, wherein it acts as a cofactor of *sod*, inhibits NADPH oxidase and stabilizes the membranes (12). However, exposure to this trace element did not influence *R. arrhizus* sensitivity to osmotic stress or modify the expression of osmotic-stress sensors *sho, ste* and p38/High Osmolarity Glycerol (HOG1) mitogen-activated protein kinase. The susceptibility to amphotericin B and posaconazole, the two main-line drugs recommended against mucormycosis (1, 3), also remained uninfluenced by zinc supplementation. Zinc uptake transporters were expectedly down-regulated under conditions of zinc enrichment. The expression of aspartic-type and serine-type proteases was also reduced, possibly owing to the observed modulation of amino-acid metabolism by this heavy metal.

Notably, the expression of some of the key virulence factors of *R. arrhizus* was enhanced by zinc, including the genes involved in iron acquisition (*ftr*1p, permease; *fre* and *fet*, reductive iron assimilation), likely attributable to the Zn-Fe homeostasis crosstalk (34). Both Ftr and Fet have been reported to be crucial for the infection process of *R. arrhizus*, as iron assimilation plays a central role in mucoralean virulence and elevated iron availability correlates with the pathogenicity of these fungi (35). Also, the expression of mucoricin, a ricin-like toxin produced by *Mucorales* (36), was up-regulated under zinc replete conditions. Mucoricin is a protein synthesis inhibitor that has been shown to promote host cell damage and vascular leakage, contributing to infection lethality (36). Elevated expression of this toxin during zinc-enriched conditions underscores the potential role played by zinc over-availability in supporting mucoralean virulence.

Taken together, the results of this study demonstrate that zinc enrichment supports *R. arrhizus* growth, and partly alleviates the effect of cell-wall and oxidative stress. Zinc-supplemented *R. arrhizus* cultures over-express multiple genes involved in growth and pathogenesis, including respiratory electron transport chain, ergosterol biosynthetic pathway, chitosan production, oxidative stress response, mucoricin, high-affinity iron permease, iron transport multi-copper oxidase and ferric-reductase-like protein. Zinc also augments the growth/metabolism of other *Mucorales* tested but not that of *A. fumigatus* and *C. albicans,* the two other common invasive fungi. The findings imply that excessive consumption of zinc supplements during COVID-19 pandemic likely contributed to the emergence of mucormycosis.

## MATERIALS AND METHODS

### Fungal strains and culture conditions

*R. arrhizus* NCCPF 710004 was used as the test strain in this study. *R. arrhizus* NCCPF 710003*, R. pusillus* NCCPF 720004, *L. corymbifera* NCCPF 700002*, C. bertholletiae* MTCC 8869, *M. indicus* MTCC 3318, *M. irregularis* MTCC 10708, *C. albicans* ATCC 90028 and *A. fumigatus* NCCPF 770372 were tested where specified. These had been procured from National Culture Collection of Pathogenic Fungi (NCCPF), Postgraduate Institute of Medical Education and Research, Chandigarh, India, and Microbial Type Culture Collection (MTCC), Institute of Microbial Technology, Chandigarh, India during our previous studies (37, 38), and were preserved in 15% glycerol (v/v) at **–**70°C. The mould strains were cultured on Sabouraud dextrose agar for 4-5 days at 37℃ (30℃ for *Mucor* spp.), and the spores were harvested using normal saline (0.85% sodium chloride; for mucoralean species) or normal saline with 0.025% Tween 20 (*Aspergillus* sp.) as described previously (38, 39). *C. albicans* ATCC 90028 was cultured on Sabouraud dextrose agar for 24 h at 37℃. The experiments to determine the impact of zinc were performed in the chemically-defined culture medium RPMI-1640 broth/agar buffered with 165 mM 3-(N-Morpholino) propane sulfonic acid to the desired pH. Zinc sulfate heptahydrate (ZnSO_4_.7H_2_O) served as the source of zinc, and the effective zinc levels were set in the culture medium accordingly. The stock solutions of zinc sulfate were prepared in water, sterilized by membrane filtration, and added to RPMI-1640 at the desired concentration. Type I water (Ultra Clear^TM^ water purification system, Evoqua Water Technologies, Günzburg, Germany) was used throughout the experiments.

### Fungal radial growth assay

RPMI-1640 agar was prepared with varying zinc concentrations (5 µM, 15 µM, 45 µM and 150 µM), and buffered to different pH (5.6, 7.0, 7.4 and 7.9). An inoculum of 1 × 10^5^ *R. arrhizus* spores was then spot-inoculated in the center of these plates, followed by incubation at 37°C for 48 h (or upto 5 days for set-ups at pH 7.9) (37). Controls without zinc supplementation were also set up. The mean diameters of fungal radial growth were measured every 24 h.

### Determination of metabolic activity

*R. arrhizus* (2.7 × 10^4^ spores/ml) was incubated in RPMI-1640 broth (pH 7.0) supplemented with zinc (5 µM, 15 µM, 45 µM and 150 µM) at 37°C in the presence of resazurin dye (final concentration, 40 µM), and the absorbance was measured at 570 and 620 nm in a microplate reader (Multiskan FC photometer, Thermo Fisher Scientific Inc., Mumbai, India) after specified time intervals. Controls without zinc supplementation and media-only blanks were also run. The percentage resazurin reduction was calculated and taken as an indicator of metabolic activity (40).

### Impact of cell-wall stress

*R. arrhizus* (1 × 10^5^ spores) was spot-inoculated on RPMI-1640 agar (pH 7.0) plates containing Congo red (250 µg/ml) as the cell-wall stressor (41), without or with zinc supplementation (5 µM, 15 µM, 45 µM and 150 µM). Media-only controls without Congo red were also set. The plates were incubated at 37°C for 48 h, and radial growth diameters were determined.

### Impact of oxidative and osmotic stress

*R. arrhizus* (1 × 10^5^ spores/ml) was exposed to hydrogen peroxide (2 µM, 6 mM and 10 mM) as the oxidative stressor (32, 33) or sodium chloride (0.145 M, 0.435 M and 1.45 M) as the osmotic stressor (42) in RPMI-1640 broth (pH 7.0) at 37°C, without or with zinc supplementation (5 µM, 15 µM, 45 µM and 150 µM). Media-only controls without the respective stress inducers were set in parallel. The spore germination was determined microscopically at defined time intervals. Germination was defined as the extension of the germ tube to a length equal to one-half the diameter of the spores (37).

### Antifungal susceptibility testing

The susceptibility of *R. arrhizus* to amphotericin B and posaconazole was determined by broth microdilution according to the guidelines of Clinical and Laboratory Standards Institute (43). A standard inoculum of 2.7 × 10^4^ spores/ml (recommended range, 0.4–5 × 10^4^ spores/ml) was treated with two-fold dilution series of the drugs (0.03–16 µg/ml) in 200 μl of RPMI-1640 (pH 7.0) at 37°C for 24 h, without or with zinc supplementation (5 µM, 15 µM, 45 µM and 150 µM), and the minimum inhibitory concentration was noted (43).

### Biofilm-forming capacity

Biofilm-forming capacity of *R. arrhizus* was studied using the microtiter-plate-based assay (39). *R. arrhizus* spores were adjusted to a count of 1 × 10^5^/ml in RPMI-1640 broth (pH 7.0), without or with zinc supplementation (5 µM, 15 µM, 45 µM and 150 µM), and 200 µl of this suspension was inoculated per well in 96-well, flat-bottomed polystyrene microtiter plates. Following incubation at 37°C for 24 h, the adhered biomass was washed with normal saline, fixed with 95% ethanol, stained with 0.5% safranin and examined under an inverted microscope (Magnus INVI, Magnus Opto Systems India Pvt. Ltd., New Delhi, India). The absorbance of the bound safranin, eluted with 200 µl of 30 % glacial acetic acid, was measured at 490 nm.

### Determination of mycelial dry weight

*R. arrhizus* (1 × 10^5^ spores/ml) was cultured in 75 ml of RPMI-1640 broth (pH 7.0) at 37°C, 150 rpm for different time intervals (2 to 8 days) without or with zinc supplementation (15 µM and 45 µM). The resulting biomass was filtered through a Whatman filter paper (grade 40; diameter, 47 mm; pore size, 8 μm), dried for 24 h at 60°C, and the mycelial dry weight was determined (37).

### RNA transcriptome sequencing

*R. arrhizus* (1 × 10^6^ spores/ml) was cultured in 10 ml of RPMI-1640 broth (pH 7.0) at 37°C, 150 rpm for 6 h without or with 45 µM zinc. The resulting growth was collected, washed and resuspended in 500 μl of RNA*later* stabilization solution for RNA extraction. Illumina-stranded RNA library preparation using NEBNext Ultra II Directional RNA library Prep Kit, and sequencing were performed by Neuberg Diagnostics Pvt. Ltd., Ahmedabad, India. The raw fastq reads were preprocessed using Fastp v.0.23.4, and the trimmed fastp reads were aligned against the SILVA database to filter the rRNA reads using RiboDetector v0.3.1. The rRNA filtered reads were aligned to the STAR indexed *Rhizopus delemar* RA 99-880 genome using STAR aligner v2.7.9a. The sorted BAM files generated from alignment were subjected to RSeQC (rseqc V4.0.0) for read distribution over genome structure and visualization using MultiQC. The rRNA and tRNA features were removed from the GTF file of the *Rhizopus delemar*, and the genome was indexed using RSEM v2.14.0. The BAM files obtained from STAR alignment were used for quantification of transcripts. edgeR annotated file was filtered based on adjusted *p*-value (false discovery rate, FDR) ≤ 0.05 and LogFoldChange ≥ 2 for differential expression. The protein sequences were extracted and subjected to eggNOG-mapper for functional annotation, BlastKOALA for KEGG annotation, and PANNZER2 for gene ontology (GO) annotation.

### Field-emission scanning electron microscopy (FE-SEM)

*R. arrhizus* (1 × 10^6^ spores/ml) was cultured in RPMI-1640 broth (pH 7.0) at 37°C, 150 rpm for 6 h without or with 45 µM zinc. The resulting suspensions were fixed with 2.5% glutaraldehyde (pH 7.2) at 4°C for 18 h, washed with Sorensen’s phosphate buffer (0.1 M, pH 7.2) plus 7% sucrose, and dehydrated in ethanol (50, 70, 80, 90 and 100% for 10, 10, 15, 15 and 20 min respectively) (38). The samples were sputter-coated with platinum and observed by FE-SEM (SU 8010, Hitachi, Japan).

### Effect of zinc on other fungal strains

The effect of zinc on growth of *R. arrhizus* NCCPF 710003*, R. pusillus* NCCPF 720004, *L. corymbifera* NCCPF 700002*, C. bertholletiae* MTCC 8869, *M. indicus* MTCC 3318, *M. irregularis* MTCC 10708 and *A. fumigatus* NCCPF 770372 was determined by radial-growth based plate assay at 37°C (30℃ for *Mucor* spp.) as described above. The growth of *C. albicans* ATCC 90028 was evaluated by determining the growth curves in RPMI-1640 containing varying zinc concentrations. Briefly, *C. albicans* was inoculated at O.D._600_ = 0.1 in RPMI-1640 buffered to different pH (5.6, 7.0, 7.4 and 7.9), without or with zinc (15 µM, 45 µM and 150 µM). The O.D._600_ measured after incubation at 37°C for 0, 2, 4, 8, 24, 48 and 72 h. Metabolic activity of all the strains was determined by resazurin dye reduction test as described above, with incubation at 37°C (30℃ for *Mucor* spp.) for a period of 6 h for *R. arrhizus, C. bertholletiae*, *M. indicus*; 12 h for *L. corymbifera*; and 24 h for *R. pusillus*, *M. irregularis, A. fumigatus* and *C. albicans* based on the dye-reduction kinetics of different strains.

### Statistical analyses

All experiments were done in triplicate and performed thrice to confirm the observations. Statistical significance was determined by Analysis of Variance test with Tukey’s multiple comparison.

## Acknowledgement

The study was supported by research funding from the Panjab University development fund. The support from the Indian Council of Medical Research (ICMR), Govt. of India under extramural research grant scheme (No. 67/2/2020-DDI/BMS) granted to RS is also acknowledged. We thank Sophisticated Analytical Instrumentation Facility (SAIF), Panjab University, Chandigarh, India for providing the FE-SEM facility.

## Supplementary data

**S1.** Normalized counts of *Rhizopus arrhizus* NCCPF 710004 gene expression after culturing in RPMI-1640 (pH 7.0) supplemented with 45 µM zinc compared with the RPMI-1640 only control, as determined by RNA transcriptome sequencing.

**S2.** Annotated file of the genes expressed by *Rhizopus arrhizus* NCCPF 710004, as determined by RNA transcriptome sequencing.

